# Amplification-Driven S100A11 Overexpression in Hepatocellular Carcinoma Is Linked to Metabolic Reprogramming, ECM Remodelling, and Immune Evasion: A Pan-Cancer Genomic Study

**DOI:** 10.64898/2026.06.15.732272

**Authors:** Stuart Lutimba, Eiman Aleem

## Abstract

**Background:** S100A11, a calcium-binding S100 family protein, is increasingly implicated in carcinogenesis, yet its molecular regulation and clinical relevance across cancers remain unclear. Hepatocellular Carcinoma (HCC) carries a dismal prognosis, in part due to a lack of reliable biomarkers for early detection and risk stratification.

**Methods:** We conducted a pan-cancer analysis of S100A11 genomic alterations across 43 studies (112,646 samples), encompassing copy number alterations, somatic mutations, and DNA methylation. HCC-specific analyses evaluated S100A11 expression, diagnostic performance, co-expression networks, and pathway enrichment using TCGA-LIHC data, with univariate and multivariate Cox regression to assess survival associations.

**Results:** S100A11 alterations were predominantly driven by copy number amplification, with the highest frequencies in lung, uterine, and hepatobiliary cancers. Copy number amplification showed a consistent inverse relationship with promoter methylation, indicating amplification-driven transcriptional activation. In HCC, S100A11 was markedly overexpressed compared with normal liver tissue, with strong diagnostic discriminatory capacity. Although high S100A11 expression trended towards inferior overall survival, this did not reach statistical significance in multivariate analysis. Co-expression and pathway analyses linked S100A11 to metabolic reprogramming, extracellular matrix remodelling, and immune dysregulation.

**Conclusion:** These findings establish S100A11 as a context-dependent oncogenic regulator highly expressed and a candidate diagnostic marker in HCC.

**Simple Summary:** S100A11, a calcium-binding S100 family protein, is increasingly implicated in carcinogenesis, yet its molecular regulation and clinical relevance across cancers remain unclear. Hepatocellular carcinoma (HCC) has a dismal prognosis, in part due to delayed detection and less effective risk stratification. In the present study, we integrated genomic, epigenetic, and transcriptomic data to characterize S100A11 alterations in HCC and assess their association with patient survival, with the aim of defining its potential as a prognostic biomarker.

**Graphical Abstract:** 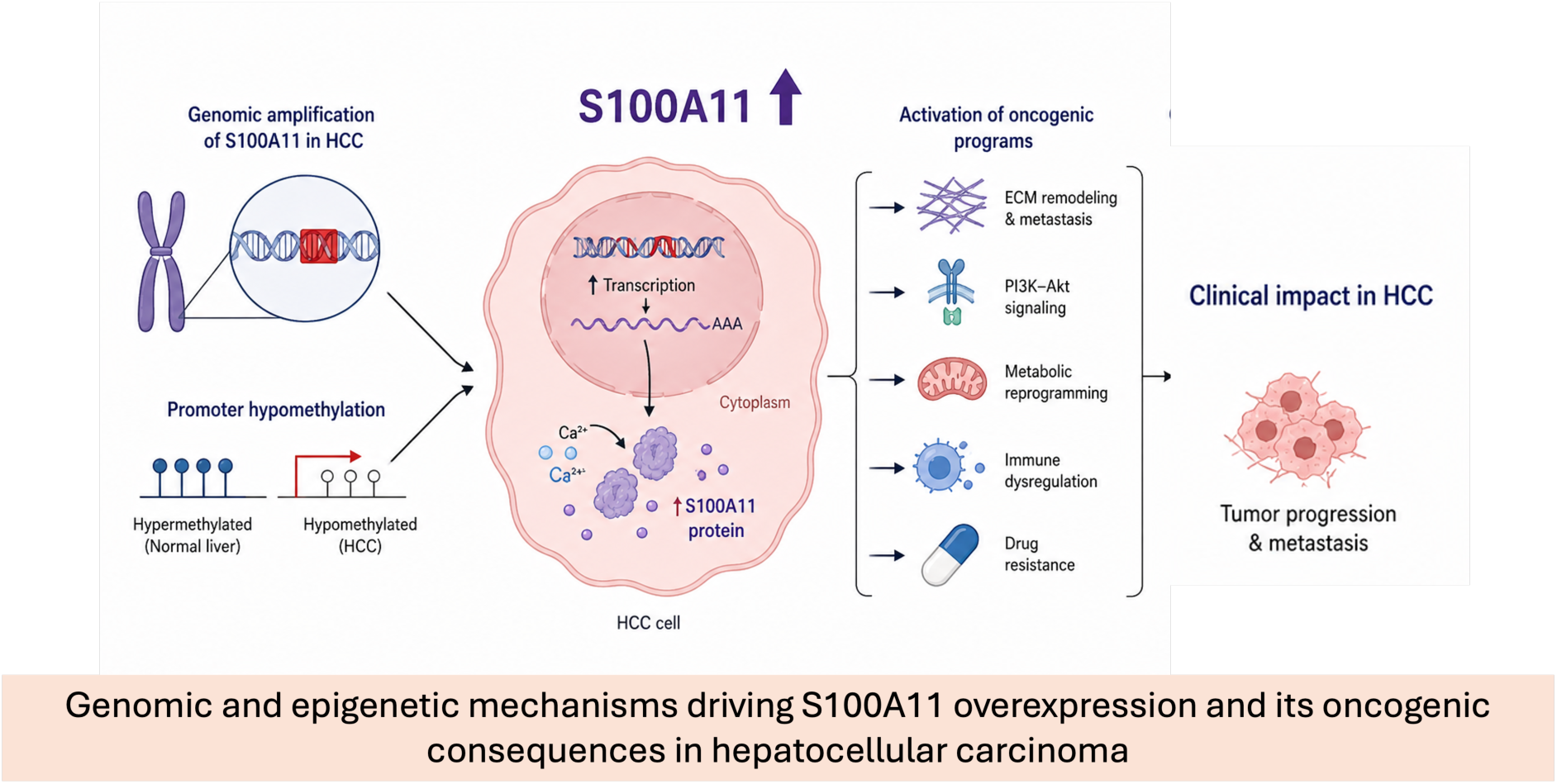

## 1. Introduction

S100A11 is a member of the S100 family of small calcium-binding proteins, which play key roles in calcium-dependent signal transduction. Members of this family regulate diverse cellular processes, including cytoskeletal organization, cell cycle progression, differentiation, migration, and stress responses, many of which are frequently dysregulated in cancer [1]. The S100 protein family comprises 25 proteins widely expressed in various cell types and tissues [2]. S100A11 has been implicated in membrane repair, actin remodelling, and transcriptional regulation, highlighting its potential role in tumourigenesis [3]. Emerging evidence has also highlighted a role for S100A11 in regulating focal adhesion dynamics [4].

Increasing evidence suggests that S100A11 is aberrantly expressed in a wide range of malignancies and contributes to cancer progression in a context-dependent manner. Elevated S100A11 expression has been reported in pancreatic, colorectal, breast, and liver cancers, where it has been associated with enhanced proliferation, epithelial–mesenchymal transition (EMT), invasion, and metastatic capacity [3,5-8]. Mechanistically, S100A11 has been shown to modulate oncogenic pathways such as TGF-β, PI3K/AKT, and MAPK signalling, thereby promoting tumour aggressiveness and resistance to therapy [6,8]. However, despite these findings, the extent to which S100A11 acts as a shared oncogenic driver versus its cancer-specific function remains unclear.

Pan-cancer genomic and transcriptomic analyses provide a powerful framework to address this question by enabling systematic comparison of genomic alterations and gene expression patterns across multiple cancer types. Large-scale datasets generated by The Cancer Genome Atlas (TCGA) have revealed both common and cancer-specific features, improving our understanding of tumour heterogeneity and identifying novel therapeutic vulnerabilities [9]. Applying a pan-cancer approach to S100A11 allows for evaluation of its genomic variations and transcriptional dysregulation across different types of cancer, its association with key oncogenic pathways, and its potential prognostic relevance.

Our pan-cancer analysis of S100A11 genetic alterations identified lung adenocarcinoma (LAC), uterine cancers and hepatobiliary cancers as exhibiting the highest frequencies of copy number alterations. As S100A11 has been previously well characterised in lung cancers [10,11,12], this study subsequently focused on a detailed transcriptional analysis of S100A11 in hepatocellular carcinoma (HCC).

HCC is the most common primary liver cancer and represents a major global health burden, ranking among the leading causes of cancer-related mortality worldwide [13]. Despite advances in targeted therapies and immunotherapy, clinical outcomes for patients with advanced HCC remain poor, largely due to extensive molecular heterogeneity and limited biomarkers for patient stratification, as well as relatively late detection of the disease. Comprehensive genomic and transcriptomic profiling has demonstrated that HCC is characterised by recurrent alterations in pathways regulating cell cycle control, WNT/β-catenin signalling, telomere maintenance, oxidative stress, and epigenetic regulation (reviewed in [14]); [15]. Transcriptome-based classifications have further identified biologically distinct HCC subtypes associated with proliferation, immune activity, and metabolic reprogramming, each with prognostic and therapeutic implications [16]. Within this complex molecular landscape, dysregulation of calcium-binding proteins and stress-response pathways has emerged as an important contributor to hepatocarcinogenesis.

Notably, increased S100A11 expression has been reported in HCC tissues compared with adjacent non-tumour liver and has been associated with poor differentiation, vascular invasion, and unfavourable clinical outcomes [8,17]. However, existing studies are largely limited to functional models or individual cohorts, and a comprehensive transcriptomic characterisation of S100A11 in HCC within a broader pan-cancer context is limited. A focused analysis of S100A11 expression and its molecular associations in HCC is therefore critically needed to clarify its biological relevance, identify co-expressed gene networks and pathways, and assess its potential as a prognostic biomarker or therapeutic target.

In this study, we conducted a pan-cancer genomic analysis of S100A11 using publicly available datasets and focused our transcriptional analysis on HCC. By integrating genomic alterations, differential expression, survival analyses, and pathway enrichment approaches, we aimed to elucidate the cancer-specific features associated with S100A11 and to define its potential prognostic and diagnostic role in HCC.

## 2. Materials and Methods

### 2.1. Multi-cohort genomic alteration profiling of S100A11

A comprehensive analysis of S100A11 genomic alterations was conducted using cBioPortal, employing a pan-cancer analysis of 43 cancer studies comprising over 112,000 samples (https://www.cbioportal.org/). The study assessed multiple types of genomic alterations, including copy number variations (amplifications, gains, shallow deletions, and deep deletions) and somatic point mutations. Data sources for these analyses included TCGA (https://www.cancer.gov/ccg/research/genome-sequencing/tcga) and complementary genomic databases integrated through cBioPortal.

DNA methylation, mutational signatures and clonality associated with S100A11 were comprehensively analyzed using TCGA datasets. Promoter methylation levels at the cg12052258 probe were extracted from Illumina HM27 and HM450 platforms and quantified as β-values, with samples stratified by copy number and mutation status to evaluate epigenetic regulation in relation to genomic alterations. DNA methylation levels are quantified as β-values derived from methylation array intensity measurements, where β represents the ratio of methylated to total signal (methylated + unmethylated). β-values range from 0.0 to 1.0, with β < 0.2 conventionally defined as hypomethylated, 0.2–0.8 as intermediate methylation, and β > 0.8 as hypermethylated. This normalization approach enables direct comparison of methylation status across samples and platforms.

Mutational processes were assessed using doublet base substitution signature contribution scores, while variant allele frequency (VAF) was used to infer clonality of S100A11 missense mutations and loss of heterozygosity in diploid samples.

### 2.2. Transcriptomic analysis of S100A11 in the TCGA-LIHC cohort

RNA sequencing data and corresponding clinical information for HCC patients were obtained from The Cancer Genome Atlas Liver Hepatocellular Carcinoma (TCGA-LIHC) project through the Genomic Data Commons (GDC) Data Portal (https://portal.gdc.cancer.gov/). The dataset comprised 424 samples, including 371 primary tumour specimens and 50 matched normal liver tissues. Gene expression data were generated using the STAR (Spliced Transcripts Alignment to a Reference) workflow, with raw counts normalized and downloaded via the TCGAbiolinks R package (version 2.28.0).

Raw RNA-seq count data were log2-transformed after adding a pseudocount of 1 to avoid undefined logarithmic values [log2(counts + 1)]. Samples were classified as tumour or normal tissue based on TCGA sample type definitions. For survival analyses, patients were dichotomized into high and low S100A11 expression groups using the median expression value as the cutoff, a standard approach in biomarker studies to ensure balanced group sizes.

### 2.3. Co-expression network and functional enrichment analysis of S100A11-associated genes in hepatocellular carcinoma

Differentially expressed genes (DEGs) associated with S100A11 were identified by comparing S100A11-high versus S100A11-low HCC tumour samples using the limma R package (version 3.56.2). RNA-seq counts were normalized via voom transformation, followed by linear modeling and empirical Bayes moderation. DEGs were defined by false discovery rate (FDR)-adjusted *p*-value *< 0.05* and log2 fold-change > 1. The top 50 DEGs (ranked by adjusted *p*-value) were visualized in heatmaps.

Co-expression analysis involved computing Pearson correlation coefficients between S100A11 and all other genes in tumour samples. The top 50 genes with strongest correlation (|r| > 0.5, *p* < 0.001) were selected for functional interrogation. Hierarchical clustering (Pearson correlation distance, complete linkage) was applied to delineate co-expression modules.

Gene Ontology (GO) and Kyoto Encyclopedia of Genes and Genomes (KEGG) pathway enrichment analyses were performed using clusterProfiler (version 4.8.2), with gene symbol-to-Entrez ID mapping via org.Hs.eg.db. Significantly enriched terms (adjusted *p* < 0.05, Benjamini-Hochberg) were ranked by gene ratio; the top 20 were visualized.

### 2.4. Evaluation of S100A11 as a prognostic and diagnostic biomarker in hepatocellular carcinoma

Overall survival (OS) was assessed using Kaplan-Meier curves generated with the survival R package, stratifying patients by S100A11 expression (high vs. low; median cutoff) and comparing groups via log-rank test (*p* < 0.05). Number-at-risk tables were included.

Prognostic impact of S100A11 genomic alterations (altered vs. unaltered; n=531) was examined using Kaplan-Meier curves and log-rank tests. Univariate and multivariate Cox proportional hazards models evaluated S100A11 expression (continuous, log2-transformed) as an independent prognostic factor, adjusting for age, sex, and tumour stage. Proportional hazards assumption was confirmed via Schoenfeld residuals. Hazard ratios (95% CI) were visualized in forest plots.

Diagnostic performance of S100A11 expression in distinguishing HCC from normal tissue was assessed by ROC analysis (pROC package), reporting area under the curve (AUC) with 95% CI (DeLong’s method), sensitivity, and specificity at Youden’s optimal cutoff.

### 2.5. Data analysis and visualization

All analyses were conducted in R (version 4.3.2). Visualizations employed ggplot2, ComplexHeatmap, pheatmap, survminer, and cowplot packages. All images were saved in TIFF (LZW-compressed) and PNG formats. Heatmaps displayed row Z-score-normalized expression with clinical annotations. Genomic alteration plots (OncoPrint, copy number, methylation) used standardized color schemes and transparency for clarity.

Statistical rigor included two-sided tests, FDR correction, complete-case analysis for missing data, and explicit reporting of sample sizes. Post-hoc power calculations confirmed adequate power (>80%) for primary analyses, with cautious interpretation for underpowered subgroups. Non-parametric (Spearman) and categorical (Fisher’s exact) tests were applied as appropriate.

## 3. Results

### 3.1. Pan-Cancer Genomic Landscape and Tissue-Specific Alterations of S100A11 Across 43 Cancer Studies

To systematically characterise the genomic alteration landscape of S100A11 across human malignancies, we performed an analysis integrating 112,646 samples across 43 independent cancer genomics studies encompassing a broad spectrum of cancer histologies, molecular subtypes, and patient populations. This multi-cohort landscape included 34 cancer types from the Cancer Genome Atlas (TCGA) in addition to a number of independent initiatives such as studies from the Memorial Sloane Kettering Cancer Center (MSK, and the MSK-IMPACT - Integrated Mutation Profiling of Actionable Cancer Targets), AMC, INSERM, METABRIC (Molecular Taxonomy of Breast Cancer International Consortium) and the SU2C (Stand Up To Cancer) initiative. Across this pan-cancer cohort, copy number amplification emerged as the dominant mechanism of S100A11 genomic dysregulation, vastly outnumbering somatic point mutations in nearly all cancer types examined (Figure 1).

**Figure 1.**
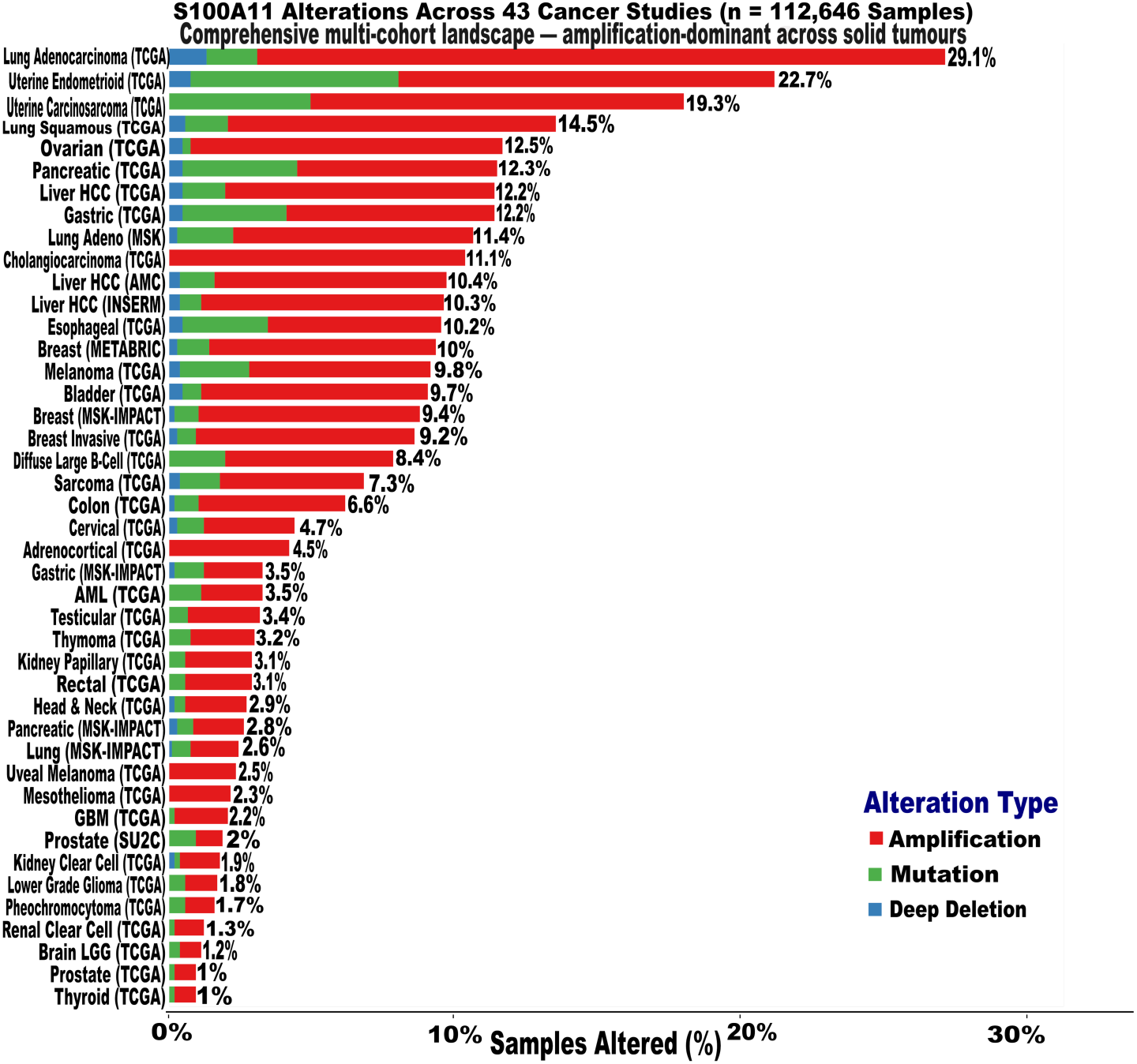
Comprehensive multi-cohort genomic alteration landscape of S100A11 across 43 cancer studies. Horizontal bar chart illustrating S100A11 alteration frequencies across 43 distinct cancer studies encompassing 112,646 samples, ranked in descending order of overall alteration burden. The x-axis represents the percentage of samples harbouring at least one S100A11 genomic alteration. Bars are colour-coded by alteration type: red, copy number amplification; green, somatic mutation; blue, deep deletion. Copy number amplification constitutes the dominant alteration class across the vast majority of tumour types represented.

The highest overall S100A11 genomic alteration frequency was observed in lung adenocarcinoma (29.1%), followed by uterine endometrioid carcinoma (22.7%), uterine carcinosarcoma (19.3%), and in lung squamous cancer (14.5%). The following four cancer types followed with an alteration frequency of 12.2% in both hepatocellular carcinoma (HCC) and gastric cancer, 12.3% in pancreatic cancer, and 12.5% in ovarian cancer, all from the TCGA cohort.

Each case driven predominantly by copy number amplification. Lung and gynecological malignancies thus represent a distinct group of cancers with a particularly elevated S100A11 genomic burden. Notably, cholangiocarcinoma (11.1%) clustered among the top ten most frequently altered cancer types in the TCGA cohort, establishing the liver as a site of recurrent S100A11 genomic activation. The HCC samples from the AMC and INSERM studies demonstrated an S100A11 genomic alteration frequency of 10.4% and 10.3%, respectively, which were also predominantly copy number amplification (Figure 1).

In contrast, somatic mutations contributed meaningfully to the alteration burden only in a subset of cancer types, most prominently uterine endometrioid and carcinosarcomas, pancreatic, gastric and esophageal carcinoma and melanoma, where mutational contributions were visually appreciable alongside amplification events. Deep deletions were rare events across the cohorts and did not constitute a primary mechanism of S100A11 dysregulation in any cancer type examined. They were more visually appreciated in lung adenocarcinoma and uterine endometroid (Figure 1).

At the lower end of the genomic alteration spectrum, thyroid carcinoma, prostate adenocarcinoma, pheochromocytoma, lower grade glioma, and kidney clear cell carcinoma each exhibited alteration frequencies at or below 2%, consistent with tissue-specific constraints on S100A11 genomic instability.

Together, these findings establish copy number amplification as the principal genomic mechanism underlying S100A11 dysregulation across human cancers, with a distinct predisposition for epithelial malignancies, and with HCC consistently occupying a place in the upper tier of alteration frequency, thus providing the primary rationale for deeper transcriptomic investigation in HCC.

### 3.2. Inverse Relationship Between S100A11 Copy Number Amplification and DNA Methylation Reveals Distinct Mechanisms of Transcriptional Dysregulation

Having established the pan-cancer prevalence of S100A11 copy number amplification, we next sought to determine whether epigenetic regulation, specifically promoter DNA methylation, operates coordinately or antagonistically with genomic copy number status to control S100A11 transcriptional activity. Interrogating methylation levels at the cg12052258 probe (Illumina HM27/HM450 merged platform) across tumour samples stratified by copy number state revealed a consistent inverse relationship between S100A11 copy number gain and promoter methylation (Figure 2A).

**Figure 2.**
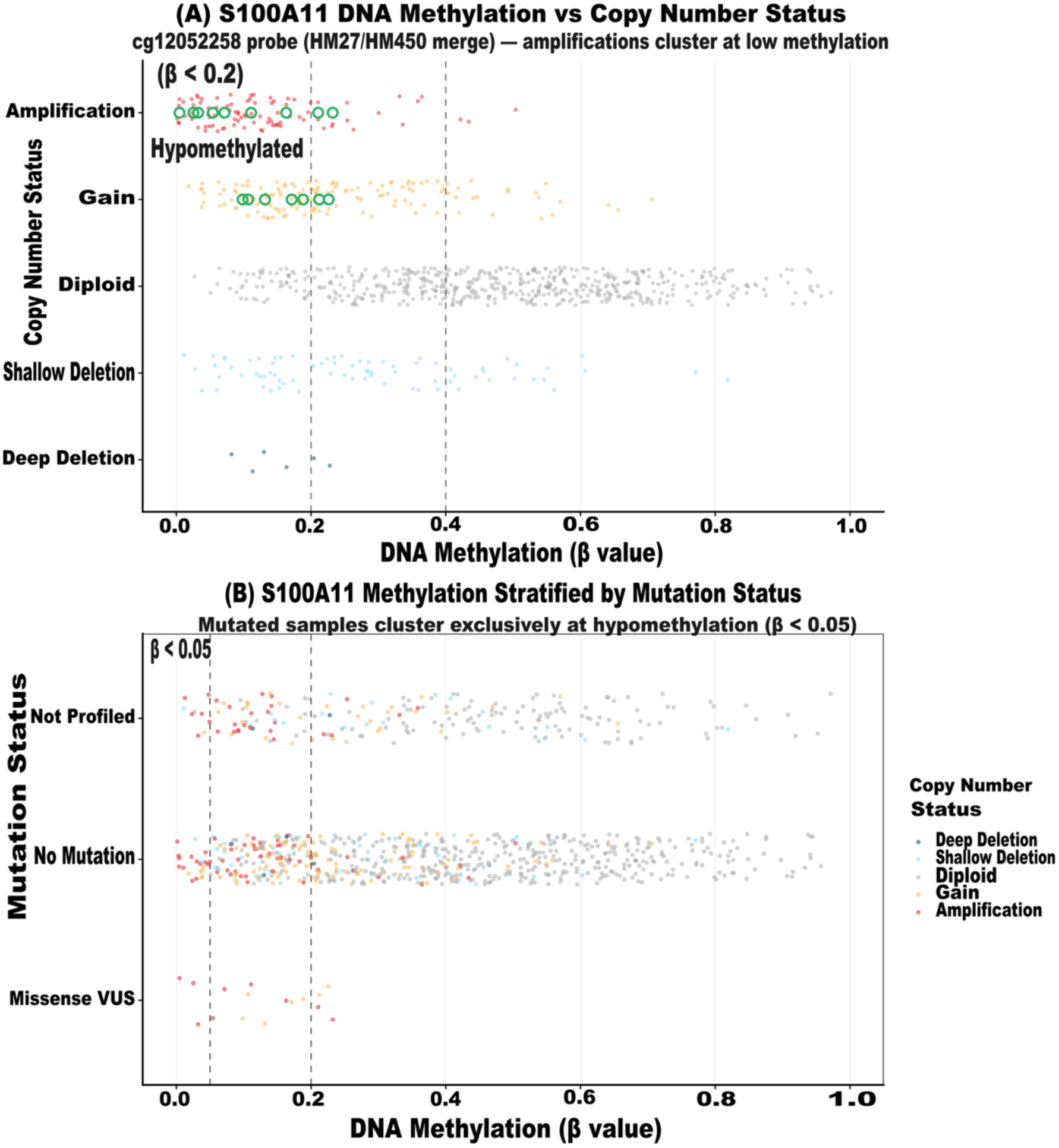
Inverse relationship between S100A11 copy number status and DNA promoter methylation across cancer samples. **(A**) Scatter plot displaying S100A11 promoter methylation levels (x-axis; cg12052258 probe, HM27/HM450 merged platform; β value range 0.0–1.0) stratified by copy number status (y-axis) across pan-cancer samples. Each data point represents an individual cancersample, colour-coded by copy number alteration class: red/pink, amplification; orange/yellow, gain; grey, diploid; light blue, shallow deletion; dark blue, deep deletion. Open green circles denote missense variants of uncertain significance (VUS). Vertical dashed lines indicate methylation thresholds at β = 0.2 and β = 0.4. Amplified and gained cancers cluster predominantly within the hypomethylated range (β < 0.2), whereas diploid cancers exhibit broad methylation variability spanning the full β range. **(B)** Scatter plot illustrating S100A11 methylation levels (x-axis; cg12052258 probe) stratified by somatic mutation status (y-axis): missense VUS (bottom), no mutation (middle), and not profiled (top). Colour coding indicates copy number status as in panel A. Missense VUS mutations occur exclusively at profound hypomethylation (β < 0.05), predominantly in diploid cancers.

Tumours harbouring copy number amplifications clustered tightly within the hypomethylated range (β < 0.2), indicating that gene dosage gain occurs preferentially in an epigenetically permissive chromatin context. Similarly, tumours with copy number gains predominantly occupied low-to-intermediate methylation values (β < 0.2–0.4), extending the pattern and collectively suggesting that amplification-driven transcriptional activation of S100A11 is facilitated by, and possibly dependent upon, concurrent promoter hypomethylation. By contrast, diploid tumours exhibited a markedly broader methylation distribution spanning the full range of β values (approximately 0.0–0.9), consistent with a model in which epigenetic regulation assumes the dominant role in controlling S100A11 expression where copy number remains unaltered. Shallow and deep deletions clustered at low-to-moderate methylation levels, though the biological significance of S100A11 in deep deletion cases is likely negligible given probable loss of functional alleles

Stratification by somatic mutation status provided a complementary and reinforcing perspective (Figure 2B). Samples harbouring S100A11 missense variants of uncertain significance (VUS) clustered exclusively at profound hypomethylation levels (β < 0.05), predominantly in the diploid copy number state, indicating that somatic mutations arise specifically within transcriptionally active, unmethylated chromatin contexts capable of supporting mutant protein expression. The no-mutation group, which comprised most samples, recapitulated the full spectrum of copy number and methylation variation observed in panel A, with amplified tumours again segregating to low methylation and diploid tumours spanning the complete methylation range. Critically, missense mutations and copy number amplifications were mutually exclusive across the cohort, with no sample co-harbouring both alteration types.

### 3.3. Characterization of the S100A11 Mutational Signatures and Clonal Architecture

To understand the functional consequences of S100A11 genomic alterations, we integrated mutational signature analysis and variant allele frequency (VAF) data to characterize the broader molecular context in which S100A11 dysregulation operates (Figure 3).

**Figure 3.**
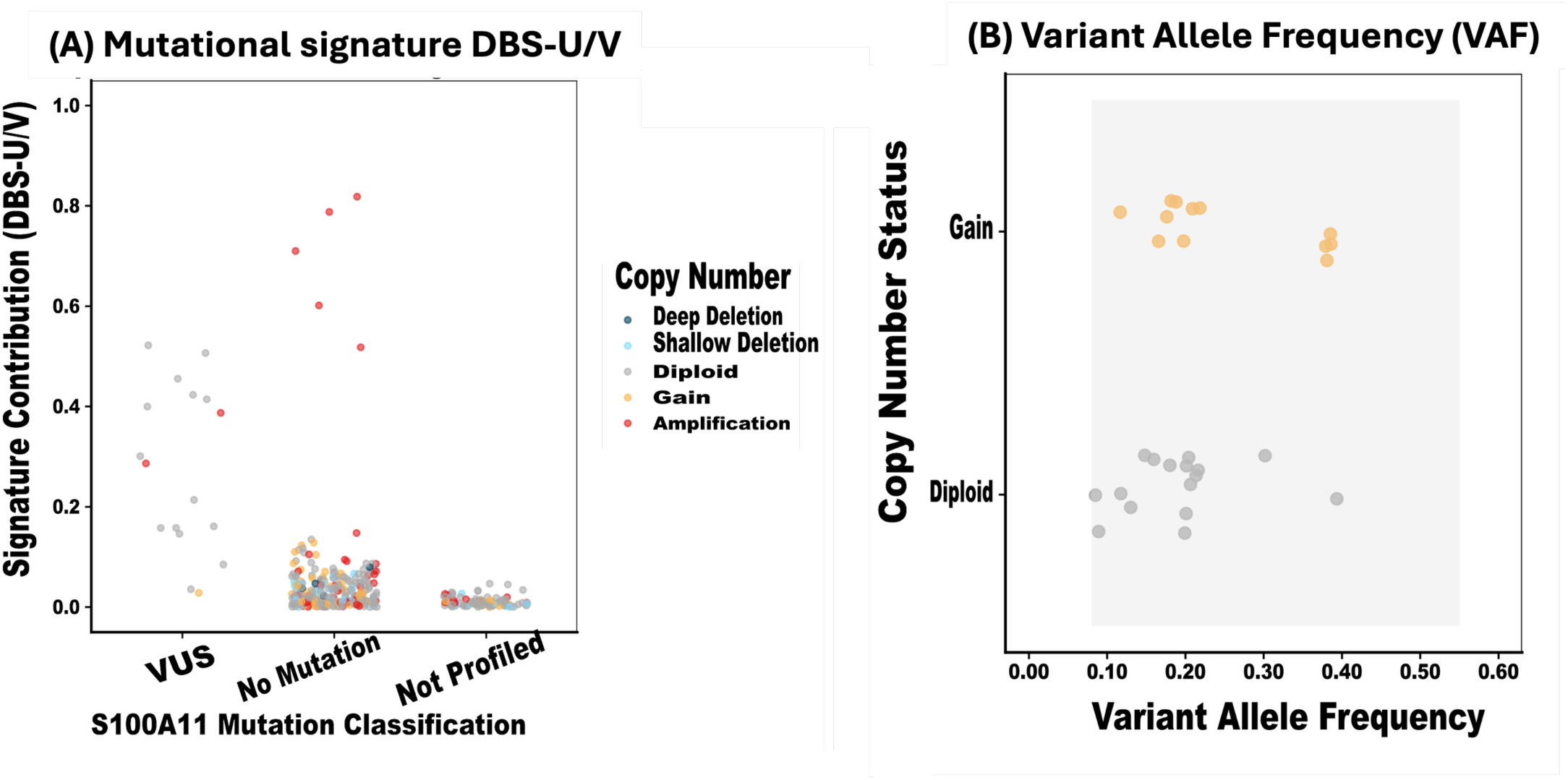
Functional characterisation of S100A11 alterations by mutational signature and clonal architecture. **(A)** Strip plot displaying doublet base substitution mutational signature contribution scores (DBS-U/V; y-axis) stratified by S100A11 mutation classification (x-axis): VUS, no mutation, and not profiled. Each data point represents an individual tumour sample, colour-coded by copy number status: red, amplification; orange/yellow, gain; grey, diploid; light blue, shallow deletion; dark blue, deep deletion. The majority of non-mutated samples cluster at DBS-U/V = 0, while a subset of amplified cases exhibits elevated signature contributions approaching 1.0. **(B)** Variant allele frequency (VAF) plot of S100A11 missense VUS mutations across copy number states (y-axis: gain, diploid). Each circle represents an individual mutation plotted by VAF (x-axis; range 0.0–0.6). Orange circles denote gain samples; grey circles denote diploid samples. The shaded region indicates the clonality assessment window.

Analysis of doublet base substitution mutational signature contributions (DBS-U/V) stratified by S100A11 mutation classification revealed that most non-mutated samples exhibited baseline signature contribution values at or near zero, consistent with absence of a dominant mutagenic process at this locus (Figure 3A). However, a subset of S100A11-amplified cases displayed markedly elevated DBS-U/V signature contributions approaching 1.0, demonstrating that copy number amplification at this locus co-occurs preferentially with broader genomic instability and elevated mutational burden.

VAF analysis of S100A11 missense mutations across copy number states provided insight into the clonal architecture and temporal sequence of mutational acquisition (Figure 3B). Missense VUS mutations displayed VAF values spanning 0.1–0.5, with gain-harbouring samples exhibiting a bimodal VAF distribution with clusters at 0.1–0.2 and approximately 0.35, consistent with the co-existence of early clonal and later subclonal mutational events within the same tumour type. Diploid samples harbouring missense mutations showed intermediate VAF values of approximately 0.15–0.2, with one additional subclonal event at approximately 0.3. The complete absence of VAF values exceeding 0.55 across all samples argues against homozygous mutation or loss-of-heterozygosity at the S100A11 locus, suggesting that retention of at least one wild-type allele confers a selective advantage.

### 3.4. S100A11 Genomic Alteration Status Does Not Independently Predict Overall Survival in Pan-Cancer TCGA Cohort

Having characterized the genomic and epigenetic landscape of S100A11 across cancer types, we next evaluated whether these alterations carry direct prognostic relevance. Kaplan-Meier survival analysis comparing patients harbouring S100A11 genomic alterations (n = 26) against unaltered controls (n = 505) across the pan-cancer TCGA cohort revealed no statistically significant difference in overall survival between groups (log-rank *p* = 0.835; HR = 1.088, 95% CI: 0.634–1.865) (Figure 4). The hazard ratio confidence interval broadly crossing unity, combined with the limited size of the altered cohort, precludes any meaningful inference regarding a survival effect attributable to S100A11 genomic alteration status alone at the pan-cancer level. Both groups exhibited broadly parallel survival trajectories throughout follow-up, and whilst transient separation of curves was observed in the intermediate follow-up window, this pattern is most parsimoniously attributable to small sample volatility in the altered group rather than a genuine biological survival difference.

**Figure 4.**
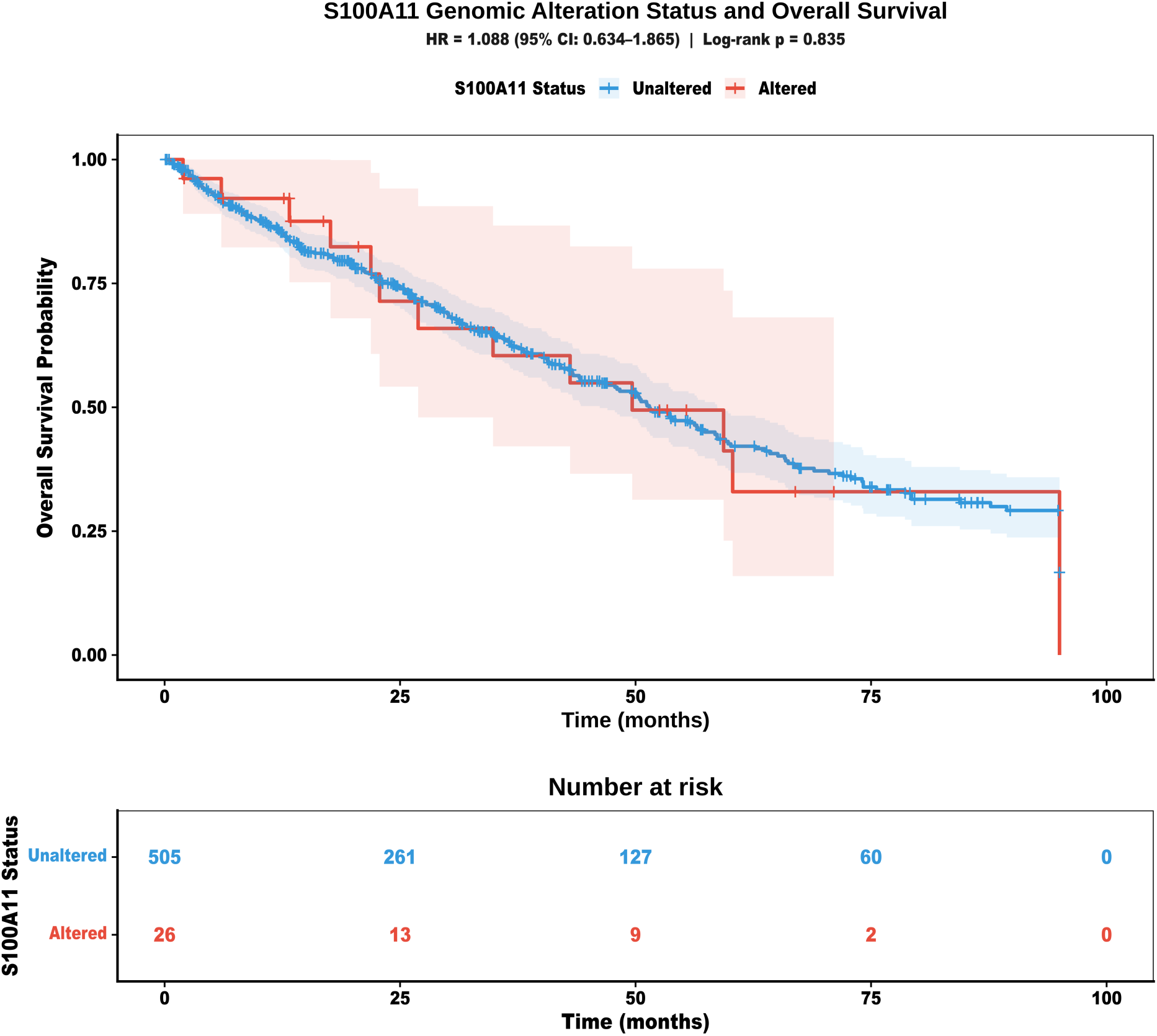
S100A11 genomic alteration status and overall survival across the pan-cancer TCGA cohort. Kaplan-Meier curves depicting overall survival probability (y-axis) over time in months (x-axis) for pan-cancer TCGA patients stratified by S100A11 genomic alteration status: unaltered (blue; n = 505) versus altered (red; n = 26). Shaded areas represent 95% confidence intervals. Tick marks indicate censored observations. The number-at-risk table below displays patients remaining under observation at specified time intervals. No statistically significant difference in overall survival was observed between the altered and unaltered groups (log-rank p = 0.835; HR = 1.088, 95% CI: 0.634–1.865).

Analysis of S100A11 transcriptional output within the HCC-specific TCGA-LIHC cohort revealed a modest, association between elevated S100A11 expression and overall survival (Figure 5). Patients dichotomized by median S100A11 expression into high (n = 186) and low (n = 185) expression groups demonstrated diverging survival trajectories, with the high-expression group exhibiting a numerically inferior survival probability across the follow-up period, though this difference did not reach conventional statistical significance in this analysis (log-rank *p* = 0.404; HR = 1.14, 95% CI: 0.84–1.54). The dashed reference line at the 50% survival threshold intersects the curves at approximately 40 months, providing a visual landmark for median survival estimation. The overlapping confidence intervals and the modest hazard ratio estimate are consistent with S100A11 expression representing one component of a broader, multivariable prognostic signature in HCC rather than a standalone determinant of survival, an interpretation that directly motivates the multivariate analyses presented in subsequent sections.

**Figure 5.**
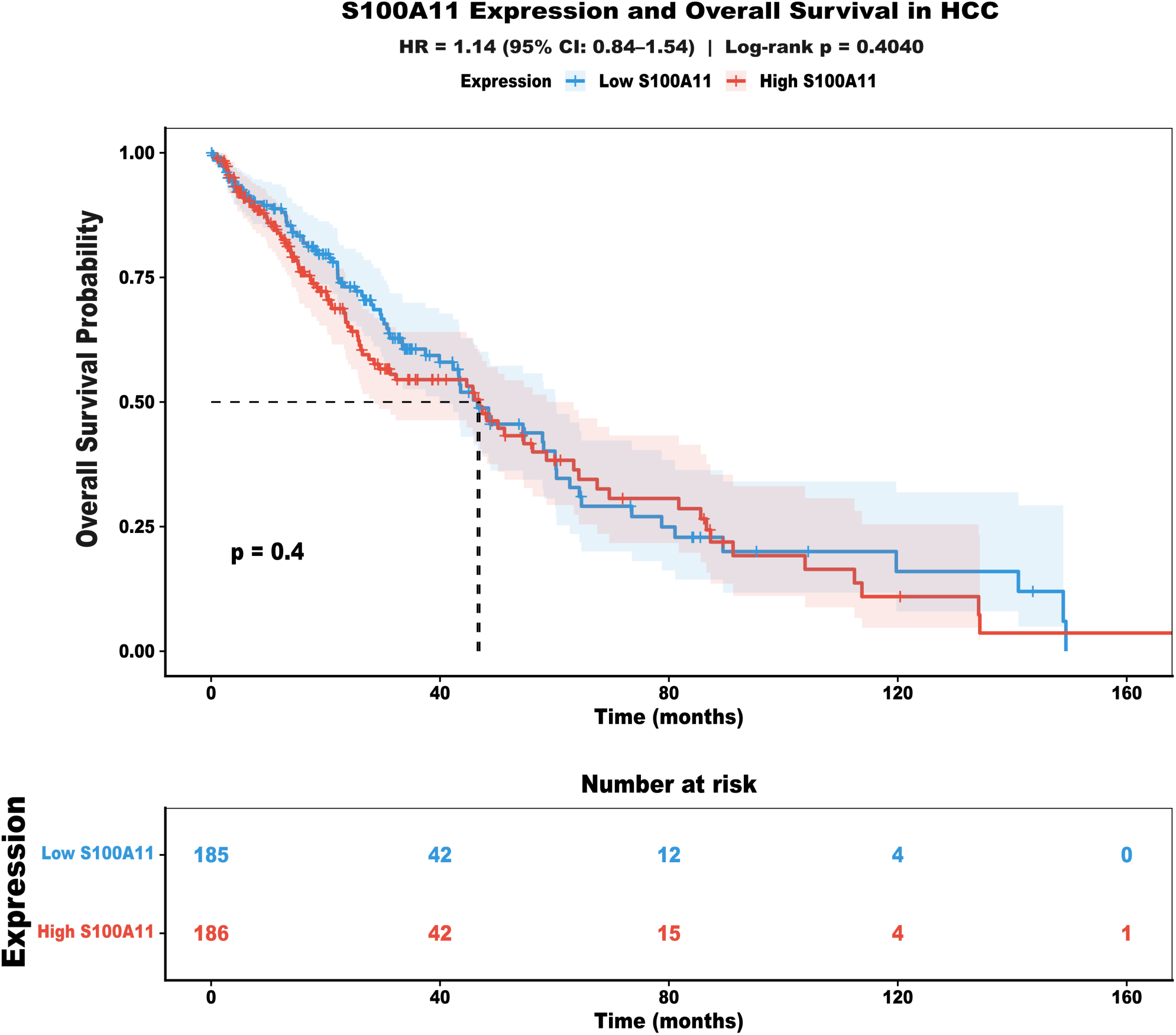
*S100A11* mRNA expression level and overall survival in hepatocellular carcinoma. Kaplan-Meier curves depicting overall survival probability (y-axis) over time in months (x-axis) for TCGA-LIHC hepatocellular carcinoma patients stratified by *S100A11* mRNA expression level: low expression (blue; n = 185) versus high expression (red; n = 186), dichotomized at the cohort median. Shaded areas represent 95% confidence intervals. Tick marks indicate censored observations. The horizontal dashed line indicates the 50% overall survival probability threshold, with the vertical dashed line marking the approximate median survival intersection at ∼40 months. The number-at-risk table below displays patients remaining under observation at specified time intervals.

### 3.5. S100A11 is Markedly Overexpressed in *HCC*

Having established the genomic and epigenetic basis of S100A11 dysregulation, we next characterised its transcriptional output directly within HCC. Comparative analysis of S100A11 mRNA expression between normal liver tissue and HCC tumour samples from the TCGA-LIHC cohort revealed highly significant upregulation of S100A11 in tumour tissue relative to normal liver parenchyma (Wilcoxon rank-sum test, p = 1.28 × 10⁻¹⁹; Figure 6A). Across 371 HCC tumour samples, S100A11 expression was markedly and consistently elevated compared with 50 normal liver tissue specimens, with the tumour distribution exhibiting both a substantially higher median expression level and a considerably broader dynamic range, reflecting transcriptional heterogeneity across the HCC cohort. The narrow, low-centred distribution of S100A11 expression in normal liver tissue contrasted sharply with the expansive, high-centred tumour distribution, underscoring the magnitude and consistency of S100A11 upregulation as a hallmark of hepatocarcinogenesis. The strength of this differential expression, spanning multiple log₂ units at the median and achieving genome-wide significance, establishes S100A11 as a robustly and reproducibly overexpressed gene in HCC, providing a strong transcriptional rationale for its evaluation as a candidate diagnostic marker.

**Figure 6.**
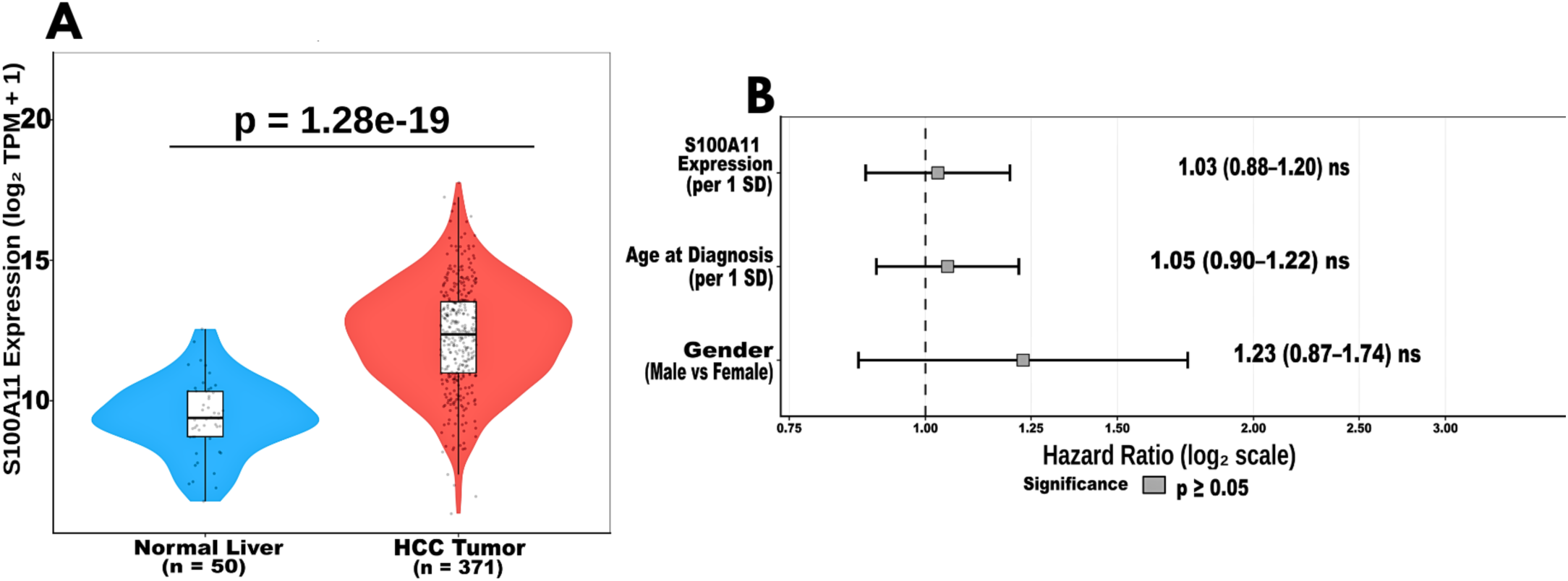
S100A11 mRNA overexpression in HCC and multivariate prognostic analysis. **(A)** Violin plots with overlaid box plots displaying S100A11 mRNA expression levels (log₂ TPM + 1; y-axis) in normal liver tissue (blue; n = 50) and HCC tumour samples (red; n = 371) from the TCGA-LIHC cohort. Individual data points are shown with jitter overlay for visualisation. The centre line of each box plot indicates the median expression value; box boundaries represent the interquartile range. Statistical significance was assessed by Wilcoxon rank-sum test (p = 1.28 × 10⁻¹⁹). **(B)** Forest plot displaying hazard ratios (HR) and 95% confidence intervals from multivariate Cox proportional hazards regression analysis of overall survival in TCGA-LIHC HCC patients with complete clinical data (n = 298). Variables included S100A11 expression (per 1 standard deviation increase; HR = 1.03, 95% CI: 0.88–1.20), age at diagnosis (per 1 standard deviation increase; HR = 1.05, 95% CI: 0.90–1.22), and gender (male versus female reference; HR = 1.23, 95% CI: 0.87–1.74). The vertical dashed line at HR = 1.0 represents the null effect. All variables are denoted as non-significant (p ≥ 0.05), indicated by grey squares. Hazard ratios are displayed on a log₂ scale.

To determine whether S100A11 expression carries independent prognostic relevance beyond its diagnostic utility, we performed multivariate Cox proportional hazards regression incorporating S100A11 expression alongside age at diagnosis and sex in 298 HCC patients with complete clinical data (Figure 6B). S100A11 expression per standard deviation increase yielded a hazard ratio of 1.03 (95% CI: 0.88–1.20, p ≥ 0.05), whilst age at diagnosis contributed an HR of 1.05 (95% CI: 0.90–1.22, p ≥ 0.05) and male sex an HR of 1.23 (95% CI: 0.87–1.74, p ≥ 0.05). None of the covariates reached conventional statistical significance. Nevertheless, the consistent directionality of all hazard ratios above unity — with S100A11 expression, older age, and male sex each trending towards increased mortality risk — is biologically coherent and consistent with established HCC epidemiology. The wide confidence intervals and modest sample size with complete covariate data (n = 298) limit statistical power, and these trends should be interpreted cautiously pending validation in larger, independent cohorts.

### 3.6. Pathway Enrichment Analysis Reveals S100A11-Associated Transcriptional Programs Governing Tumour Invasion, Metabolic Reprogramming, and Immune Regulation in HCC

To delineate the broader biological programs associated with S100A11 transcriptional upregulation in HCC, we performed KEGG pathway and Gene Ontology Biological Process (GO-BP) enrichment analyses on differentially expressed genes between S100A11-high and S100A11-low tumour groups. KEGG pathway analysis identified focal adhesion as the most significantly enriched pathway in S100A11-high tumours (gene ratio ∼5.5%, FDR < 0.05), followed by ECM–receptor interaction and PI3K–Akt signalling (Figure 7A).

**Figure 7.**
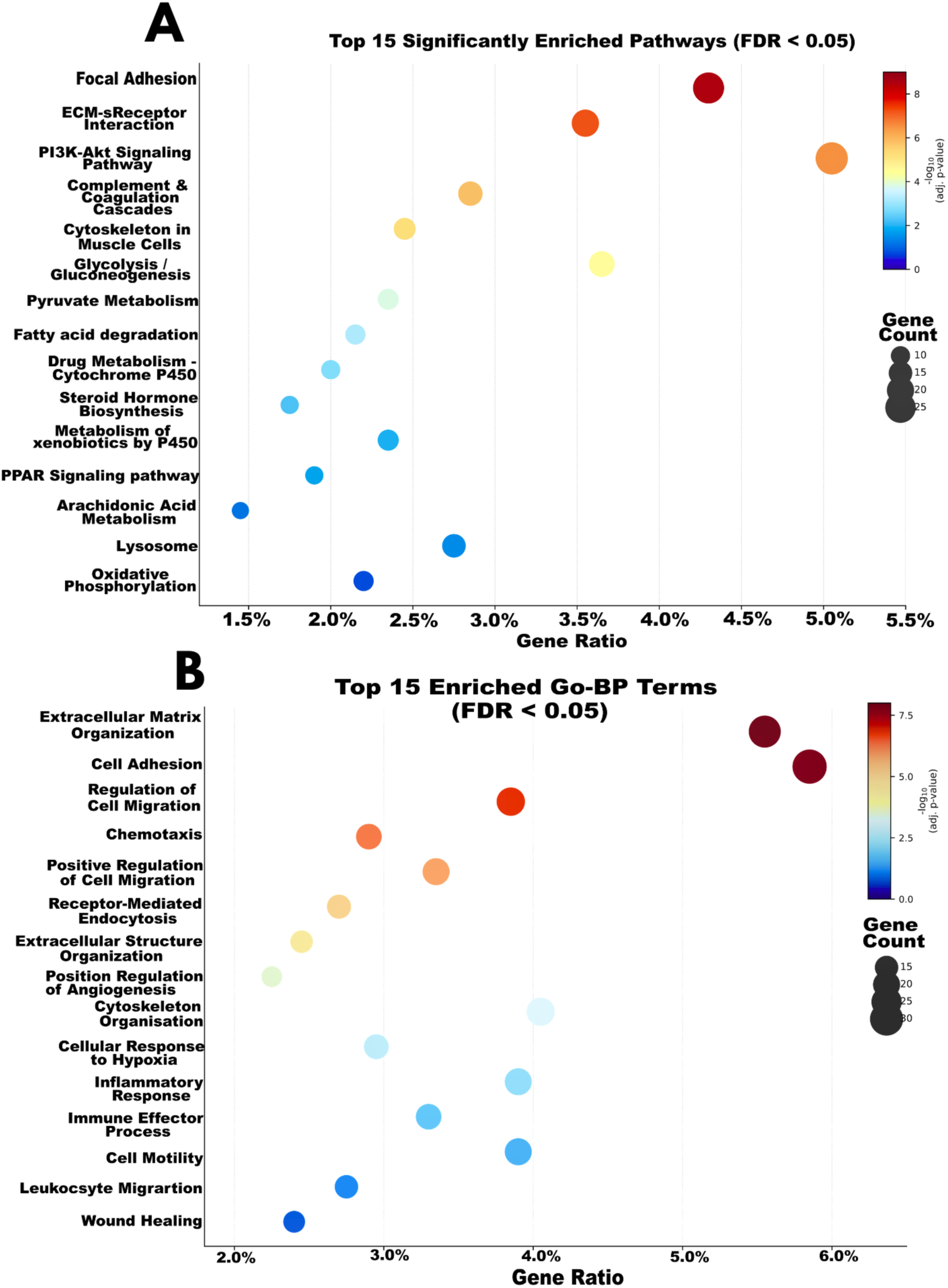
Pathway enrichment analysis in S100A11-high hepatocellular carcinoma. Dot plots displaying the top 15 significantly enriched terms (FDR < 0.05) identified from differentially expressed genes between S100A11-high and S100A11-low HCC tumours in the TCGA-LIHC cohort. The x-axis represents the gene ratio (proportion of pathway/term genes present among differentially expressed genes). Dot size is proportional to gene count and dot colour indicates statistical significance scaled by −log₁₀(adjusted p-value): red/dark red, highest significance; blue, lower significance. **(A) KEGG pathway enrichment.** Gene count scale: 10–25. Focal adhesion was the most significantly enriched pathway (gene ratio ∼5.5%), followed by ECM-receptor interaction and PI3K-Akt signalling. Metabolic pathways including glycolysis/gluconeogenesis, fatty acid degradation, and oxidative phosphorylation were also significantly enriched, alongside drug metabolism and PPAR signalling pathways. **(B) Gene Ontology Biological Process enrichment.** Gene count scale: 15–30. Extracellular matrix organisation and cell adhesion were the most significantly enriched terms (gene ratio ∼6%), followed by regulation of cell migration, chemotaxis, and receptor-mediated endocytosis. Immune-related terms including inflammatory response, immune effector process, and leucocyte migration were also significantly enriched, implicating S100A11 overexpression in tumour immune microenvironment dysregulation.

Notably, multiple metabolic pathways were significantly enriched, including glycolysis/gluconeogenesis, pyruvate metabolism, fatty acid degradation, and oxidative phosphorylation, collectively indicating that S100A11-high HCC tumours undergo extensive metabolic reprogramming. The co-enrichment of drug metabolism pathways including cytochrome P450-mediated xenobiotic metabolism, steroid hormone biosynthesis, and PPAR signalling further suggests that elevated S100A11 expression may be associated with altered drug biotransformation capacity, with implications for therapeutic response and potentially resistance in HCC.

GO-BP enrichment analysis provided complementary and orthogonal biological resolution, with extracellular matrix organisation and cell adhesion emerging as the most significantly enriched terms (gene ratio ∼6%, FDR < 0.05), closely followed by regulation of cell migration and chemotaxis (Figure 7B). Importantly, immune-related GO-BP terms including inflammatory response, immune effector process, and leucocyte migration were significantly enriched in S100A11-high tumours, suggesting that elevated S100A11 expression is coupled to immune microenvironment dysregulation and may influence the balance between pro- and anti-tumour immune activity within the HCC tumour milieu. Additional enrichment of wound healing, cellular response to hypoxia, and positive regulation of angiogenesis collectively indicate that S100A11-high HCC occupies a transcriptionally aggressive state characterized by the simultaneous activation of invasion, immune evasion, and adaptive stress response programs.

### 3.7. Co-expression Network Analysis Identifies Distinct S100A11-Associated Transcriptional Modules Governing Tumour Invasion and Immune Remodelling in HCC

To delineate the co-expression network of S100A11 in HCC and identify co-regulated genes that may cooperate in driving tumour progression, we computed Pearson correlation coefficients between S100A11 and all expressed genes across TCGA-LIHC tumour samples only (n = 371 primary HCC tumours). We selected the top 50 genes with the strongest correlation to S100A11 (|r| > 0.5, p < 0.001) for hierarchical clustering and visualization.

The resulting correlation heatmap revealed two principal co-expression modules with distinct biological identities, delineated by hierarchical clustering into upper and lower gene clusters (Figure 8). The two identified modules represent two distinct co-regulated transcriptional programs within HCC tumours, identified through hierarchical clustering based on Pearson correlation distance, with strong positive correlations among genes within each module and weaker correlations between modules.

**Figure 8.**
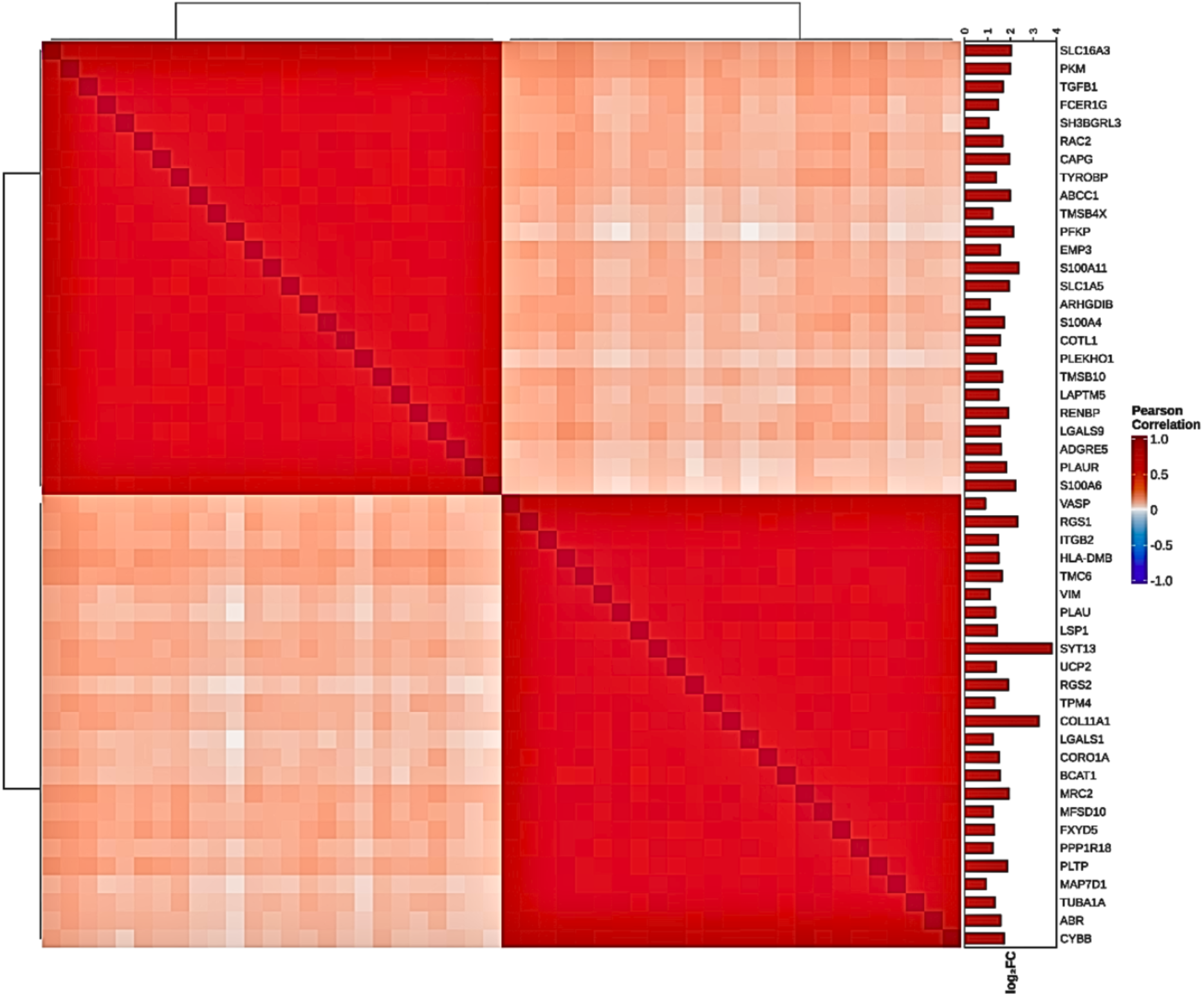
Co-expression correlation heatmap of S100A11-associated genes in hepatocellular carcinoma. Symmetric Pearson correlation heatmap displaying pairwise correlation coefficients between S100A11 and the top co-expressed genes across TCGA-LIHC HCC tumour samples. Genes are arranged by hierarchical clustering (complete linkage, Pearson distance), revealing two principal co-expression modules delineated by the dendrogram. Colour scale represents Pearson correlation coefficient: deep red, strong positive correlation (r approaching 1.0); white/light pink, weak or absent correlation (r approaching 0); blue, negative correlation. The upper module encompasses S100A11 and co-regulated genes involved in metabolic reprogramming (PKM, SLC16A3, SLC1A5), cytoskeletal organisation (TMSB4X, TMSB10, CAPG), S100 family members (S100A4, S100A6), and innate immune signalling (TYROBP, FCER1G, LGALS9, ADGRE5). The lower module comprises genes associated with actin dynamics (VASP, TPM4, CORO1A, TUBA1A), ECM remodelling (COL11A1, VIM), integrin signalling (ITGB2), and immune effector processes (HLA-DMB, CYBB, MRC2). The right sidebar displays log-fold-change values for each gene between S100A11-high and S100A11-low HCC tumours, with red bars indicating positive differential expression.

The upper co-expression module exhibited strong mutual positive correlations (Pearson r approaching 1.0), indicating tight transcriptional co-regulation. S100A11 co-clusters with other S100 family members (S100A4, S100A6), cytoskeletal regulatory proteins (TMSB4X, TMSB10, CAPG), innate immune and myeloid-associated genes (TYROBP, FCER1G, LGALS9, ADGRE5), metabolic enzymes (PKM, SLC16A3, SLC1A5), and the invasion-associated receptor PLAUR. This composition collectively positions S100A11 at the convergence of metabolic reprogramming, cytoskeletal remodelling, and innate immune dysregulation, three hallmarks of aggressive HCC. The co-clustering of TGFB1 within this module further implicates S100A11 in TGF-β-mediated epithelial–mesenchymal transition and immunosuppressive signalling, consistent with the pathway enrichment findings presented above. Notably, the myeloid gene signature within this module, anchored by TYROBP, FCER1G, and ADGRE5, suggests that S100A11-high tumours may be characterised by preferential innate immune infiltration, a pattern consistent with an immunosuppressive rather than cytotoxic tumour microenvironment.

The lower co-expression module similarly demonstrated strong intra-cluster positive correlations and was enriched for genes governing actin cytoskeleton dynamics (VASP, TPM4, CORO1A, TUBA1A), integrin-mediated adhesion (ITGB2), extracellular matrix remodelling (COL11A1, VIM), and immune effector functions (HLA-DMB, CYBB, LSP1, MRC2). Although S100A11 itself resides in the upper module, the biological programs represented in the lower module — ECM remodelling, cytoskeletal reorganisation, and immune dysregulation — are strikingly concordant with those of the upper module, indicating that these processes are robustly and independently activated across S100A11-high HCC rather than being an artefact of a single co-expression cluster.

The inter-cluster correlation was positive but attenuated relative to intra-cluster correlations, as evidenced by the lighter off-diagonal quadrants in Figure 8, indicating partial rather than complete transcriptional co-regulation between these two programs. The log-fold-change sidebar confirmed that the majority of genes within both modules were positively differentially expressed in S100A11-high tumours, underscoring the coordinated transcriptional upregulation of invasion, immune remodelling, and metabolic reprogramming programs as defining features of the aggressive HCC molecular state associated with elevated S100A11 expression.

## 4. Discussion

This study provides a multi-layered characterisation of S100A11 dysregulation across human malignancies, with particular significance in hepatocellular carcinoma (HCC). By integrating pan-cancer genomics, epigenetic profiling, and HCC-focused transcriptomics and co-expression network analyses, we demonstrate that S100A11 operates within a complex regulatory landscape in which genomic and epigenetic mechanisms converge to shape tumour behaviour, consistent with recent pan-cancer analyses identifying S100A11 as a broadly upregulated oncogenic factor associated with tumour progression and immune modulation [18].

A central finding is that copy number amplification represents the dominant mechanism of S100A11 genomic dysregulation across cancers. Previous studies have characterised S100A11 primarily through its transcriptional overexpression in individual tumour types, including breast, colorectal, and hepatocellular carcinoma [5-8]. Our results extend these observations by demonstrating that overexpression is frequently underpinned by genomic amplification, with the highest frequencies in epithelial malignancies including lung adenocarcinoma, uterine cancers, and HCC; consistent with pan-cancer analyses of the S100 protein family demonstrating lineage-dependent expression patterns and associations with immune infiltration [19]. These findings align with earlier work in the Karpas-299 anaplastic large cell lymphoma cell line, in which elevated S100A11 mRNA was attributed to DNA amplification [20].

We further identify a robust inverse relationship between S100A11 copy number amplification and promoter methylation, indicating that transcriptional activation requires an epigenetically permissive chromatin state, consistent with models of oncogene activation in which promoter hypomethylation cooperates with genomic amplification to sustain transcriptional upregulation [21,22]. The frequency of somatic mutations was very low compared with amplification, consistent with reported S100A11 mutation rates of approximately 5% in low-grade glioma [23] and across pan-cancer datasets [18]. Strikingly, amplification and somatic mutation were mutually exclusive across the entire cohort, defining two molecularly distinct routes to S100A11 dysregulation: amplification-driven overexpression in hypomethylated contexts, and mutation-driven alteration in transcriptionally active diploid tumours, each with distinct implications for patient stratification and therapeutic targeting. This mutual exclusivity reflects a broader principle in cancer genomics whereby distinct alteration types converge on the same oncogenic pathway without co-occurring, as exemplified by mutual exclusivity in PI3K, p53, and Rb pathway alterations in glioblastoma, and between BRCA1/2 inactivation and CCNE1 amplification in serous ovarian cancer [24].

From a clinical perspective, Kaplan–Meier analysis revealed no statistically significant survival difference between patients with S100A11 genomic alterations and unaltered controls. The wide confidence interval and limited size of the altered cohort preclude meaningful inference regarding a survival effect attributable to genomic alteration status alone. Transcriptional upregulation in HCC emerges as more biologically relevant, with markedly elevated expression in tumour relative to normal liver and a non-significant but directionally consistent trend towards inferior overall survival in S100A11-high patients, underscoring the importance of evaluating transcriptional rather than purely genomic endpoints for biomarker assessment [18]. These observations are consistent with reports associating S100A11 expression with poor survival in breast cancer [25], its identification as a serum biomarker in non-metastatic pancreatic cancer [26], and its protein expression as an unfavourable prognostic indicator in surgically resected pancreatic adenocarcinoma [27]. The strong discriminatory capacity of S100A11 expression in distinguishing HCC from normal liver tissue supports its candidacy as a diagnostic marker, warranting prospective validation, which is consistent with prior reports of a diagnostic role for S100A11 in HCC [8,28-30].

Mechanistically, pathway enrichment and co-expression analyses converge on a model in which S100A11 overexpression drives tumour aggressiveness through coordinated activation of ECM remodelling, cytoskeletal reorganisation, metabolic reprogramming, and immune modulation. The enrichment of focal adhesion and PI3K–Akt signalling is consistent with experimental evidence linking S100A11 to invasion and metastasis in HCC through AKT and ERK signalling [8], and emerging evidence implicates intracellular S100A11 in additional oncogenic mechanisms including desmosome-associated signalling, β-catenin/TCF activation and focal adhesion [31,4].

The enrichment of metabolic pathways including glycolysis, pyruvate metabolism, fatty acid degradation, and oxidative phosphorylation suggests S100A11 contributes to the metabolic plasticity central to tumour adaptation and therapeutic resistance [32]. HCC is characterised by extensive metabolic reprogramming encompassing glycolysis, the pentose phosphate pathway, amino acid metabolism, and fatty acid metabolism [33], and the sequential shift from gluconeogenesis and glycogen storage to aerobic glycolysis is a recognised early event in hepatocarcinogenesis [34]. Insulin receptor pathway upregulation has similarly been reported as potential early regulator of this metabolic shift [35], and the precise contribution of S100A11 to metabolic reprogramming in HCC warrants dedicated investigation.

The enrichment of cytochrome P450-mediated drug metabolism and PPAR signalling pathways raises further hypotheses regarding therapeutic resistance. Sorafenib, the first-line agent for advanced HCC, is metabolised primarily through CYP3A4-mediated oxidation, and hepatic CYP activity is a recognised determinant of pharmacokinetics and treatment response [36,37]. Sorafenib metabolism is significantly altered in HCC tumour tissue relative to normal liver, with reduced CYP expression potentially contributing to therapeutic failure [38]. The enrichment of these pathways in S100A11-high tumours raises the possibility that elevated S100A11 is associated with dysregulated drug biotransformation, thus warranting functional validation.

The enrichment of immune-related pathways is consistent with an active role for S100A11 in shaping the tumour immune microenvironment. Cancer cell-derived S100A11 promotes monocyte recruitment and macrophage infiltration in a dose-dependent manner in ER+ breast cancer models, with S100A11 silencing reducing monocyte infiltration and expression of the immunosuppressive marker CD206, establishing S100A11 as a paracrine regulator of cancer–macrophage crosstalk [39]. Consistent with this, S100A11 expression correlates with M2 macrophage polarisation and an immunosuppressive microenvironment in glioma [40], and single-cell RNA sequencing data in HCC implicate S100A11 in regulating tumour-infiltrating immune cells, particularly T-cell function via NF-κB signalling [41]. Our co-expression network reinforces this: the upper transcriptional module, in which S100A11 resides, is anchored by TYROBP, FCER1G, and ADGRE5, established transmembrane immune signalling adaptors mediating innate immune activation via ITAMs and recognised markers of macrophage identity and myeloid infiltration [42,43], thus consistent with a preferentially immunosuppressive rather than cytotoxic tumour microenvironment. Pan-cancer analyses have similarly linked S100A11 to immune checkpoint regulation and immunosuppressive cell infiltration [18], collectively positioning S100A11 as a mediator of tumour–immune crosstalk with implications for immunotherapy response in HCC.

The co-expression network identifies two partially independent transcriptional modules linking S100A11 with metabolic, cytoskeletal, and immune regulatory programs. Co-expression with S100A4, S100A6, and TGFB1 suggests coordinated activation of epithelial–mesenchymal transition, invasion, and immune evasion pathways, consistent with systems- level analyses of S100 protein function in cancer [1]. The biological programs across both moduless, ECM remodelling, cytoskeletal reorganisation, and immune dysregulation, map plausibly onto established HCC transcriptomic subclasses, including the proliferative/TGF-β-enriched and immune-active subtypes [16], a correspondence warranting direct investigation in subtype-annotated cohorts.

The present study has the following limitations: Reliance on retrospective, publicly available data introduces potential cohort biases, and the relatively small number of S100A11-altered cases limits statistical power, precluding definitive conclusions regarding the prognostic impact of genomic and transcriptomic alteration statuses. Experimental validation is required to establish causality, particularly regarding the interplay between amplification, epigenetic remodelling, and downstream signalling. The study also lacks protein-level characterisation, as mRNA expression alone does not necessarily reflect protein abundance or activity. Further studies utilizing larger HCC cohorts and independent replication in prospective or externally validated datasets is essential before clinical translation of these findings.

## 5. Conclusions

This study provides an integrated genomic and epigenetic characterisation of S100A11 dysregulation across human cancers, with a focused transcriptomic investigation in HCC. Copy number amplification, operating in concert with promoter hypomethylation, is the dominant mechanism driving S100A11 overexpression across epithelial malignancies, whilst the mutual exclusivity of amplification and somatic mutation defines two mechanistically distinct routes to dysregulation, placing S100A11 within a broader model of epigenetically permissive oncogene activation.

In HCC, S100A11 is robustly overexpressed relative to normal liver tissue with strong diagnostic discriminatory capacity, and its transcriptional network converges on tumour invasion, metabolic reprogramming, immune evasion, and potential therapeutic resistance. The myeloid-enriched co-expression signature anchored by TYROBP, FCER1G, and ADGRE5 implicates S100A11 in shaping an immunosuppressive tumour microenvironment with potential consequences for immunotherapy response. Whilst the multivariate survival analysis did not reach statistical significance, the consistent directionality of the prognostic trend supports continued investigation of S100A11 as a clinically relevant biomarker in HCC.

Future studies should prioritise protein-level validation in independent prospective cohorts, along with functional studies in HCC models to establish causality, preclinical and clinical testing of the association between S100A11 and resistance to sorafenib. Additionally, replication studies in multi-centre subtype-annotated cohorts should be conducted before clinical translation is considered. The emerging role of S100A11 in focal adhesion and metabolic reprogramming should be investigated during the transition from preneoplastic to neoplastic phenotype in HCC. Collectively, these findings establish S100A11 as a context-dependent oncogenic regulator and a candidate for both diagnostic evaluation and therapeutic targeting in HCC.

## Supplementary Materials

No additional supplementary materials are included for this article.

## Author Contributions

Conceptualization EA.; methodology SL and EA, software, bioinformatic analysis, data curation SL; validation and resources, SL and EA, writing, EA and SL.; visualization, SL.; supervision, EA.; project administration, EA. All authors have read and agreed to the published version of the manuscript.”

## Institutional Review Board Statement

Not applicable as the study didn’t involve humans or animals.

## Informed Consent Statement

Not applicable.

## Data Availability Statement

All data analysed in this study are derived from publicly available repositories. S100A11 genomic alteration data were obtained from cBioPortal (https://www.cbioportal.org/), encompassing 43 cancer studies and 112,646 samples sourced from The Cancer Genome Atlas (TCGA; https://www.cancer.gov/ccg/research/genome-sequencing/tcga) and additional independent cohorts (MSK-IMPACT, AMC, INSERM, METABRIC, and SU2C). DNA methylation and mutational signature were accessed via RNA sequencing data and clinical information for the HCC-specific analyses were obtained from the TCGA Liver Hepatocellular Carcinoma (TCGA-LIHC) project through the Genomic Data Commons (GDC) Data Portal (https://portal.gdc.cancer.gov/).

